# Cryo-electron tomography pipeline for plasma membranes

**DOI:** 10.1101/2024.06.27.600657

**Authors:** Willy W. Sun, Dennis J. Michalak, Kem A. Sochacki, Prasanthi Kunamaneni, Marco A. Alfonzo-Méndez, Andreas M. Arnold, Marie-Paule Strub, Jenny E. Hinshaw, Justin W. Taraska

## Abstract

Cryo-electron tomography (cryoET) provides sub-nanometer protein structure within the dense cellular environment. Existing sample preparation methods are insufficient at accessing the plasma membrane and its associated proteins. Here, we present a correlative cryo-electron tomography pipeline optimally suited to image large ultra-thin areas of isolated basal and apical plasma membranes. The pipeline allows for angstrom-scale structure determination with sub-tomogram averaging and employs a genetically-encodable rapid chemically-induced electron microscopy visible tag for marking specific proteins within the complex cell environment. The pipeline provides fast, efficient, distributable, low-cost sample preparation and enables targeted structural studies of identified proteins at the plasma membrane of cells.

## Introduction

Cryo-electron tomography (cryoET) offers a unique perspective of high-resolution protein structure within the crowded cellular environment. While cryoET in cells is powerful, it has two fundamental constraints. First, samples must be thinner than a typical eukaryotic cell [1]. Second, low signal to noise images and a crowded environment make it challenging to identify proteins smaller than 400 kDa [2], making it currently unsuitable for >99% of the human proteome [3]. These limitations and the need for increased efficiency and accessibility motivates the development of new cell-thinning and protein-identification methods.

Cryo-focused ion or plasma beam (FIB)-milling has become a staple for cryoET by producing thin (<200 nm) samples rich in cellular structures from the heart of cells and tissues. FIB-milling is, however, time-consuming, expensive, complex, and not well-suited for the plasma membrane. Thus, other methods are needed. Cell unroofing, a fast and inexpensive technique classically used for platinum replica electron, atomic force, and fluorescence microscopies, isolates cellular plasma membranes with a thin associated protein cortex [4, 5]. Yet, producing plasma membranes on fragile cryo-EM grids is not trivial [6, 7], and has not been used or characterized for high resolution cryoET.

Another common problem for cryoET is protein identification. To address this, EM-visible protein cages are an attractive, genetically-encodable solution. Encapsulins (20-42 nm) [8–10] and iron-sequestering ferritins (12 nm) [11, 12] are two examples of established electron microscopy tags with unique shapes. While these tags are large in comparison to the average protein or protein complex, rapamycin-induced linkages have been used to tether these oligomeric complexes to proteins inside living cells [10, 12].

However, encapsulin cages can require 15 minutes to several hours to label their targets and are more sterically inhibiting than ferritin. The rapamycin-inducible fluorescent ferritin tagging system, FerriTag, is faster, binding to its target within seconds [12] but has not been used for cryoET. While previous thin-section transmission EM studies loaded FerriTag with iron to enhance contrast [12], metal contrast in cryoET can obscure nearby proteins of interest and perturb high-resolution structural information.

Here, we present a cell unroofing workflow that generates isolated basal or apical plasma membranes for correlative cryoET. We show that, when cells are cultured on EM grids, membrane-associated protein complexes adapt to the grid surface topography. We demonstrate that cell unroofing preserves sub-nanometer resolution in ribosomes. We use correlative light and electron microscopy (CLEM) and iron-free FerriTag as a chemically-inducible tag to locate and observe clathrin associated proteins. In summary, we present and characterize a new optimized pipeline for cryoET of plasma membranes capable of identifying proteins for structural cellular biology.

## Results

### Basal and apical plasma membranes can be isolated on grids

Because of their different possible structures and biological functions, we sought to isolate basal and apical plasma membranes of cultured cells for cryoET. For basal membranes, cells were grown on Quantifoil cryoET grids. The grid was adhered to a coverslip using a PDMS (polydimethylsiloxane) stencil before cell plating. The stencil-grid-coverslip assembly could be handled as a single unit, minimizing direct grid manipulation during cell unroofing (Fig. 1a). After one day of culture, unroofing was performed with a stream of paraformaldehyde-containing buffer from a syringe, with a flow tunable by 0.7-0.8 bar compressed air (Fig. S1).

**Figure 1.**
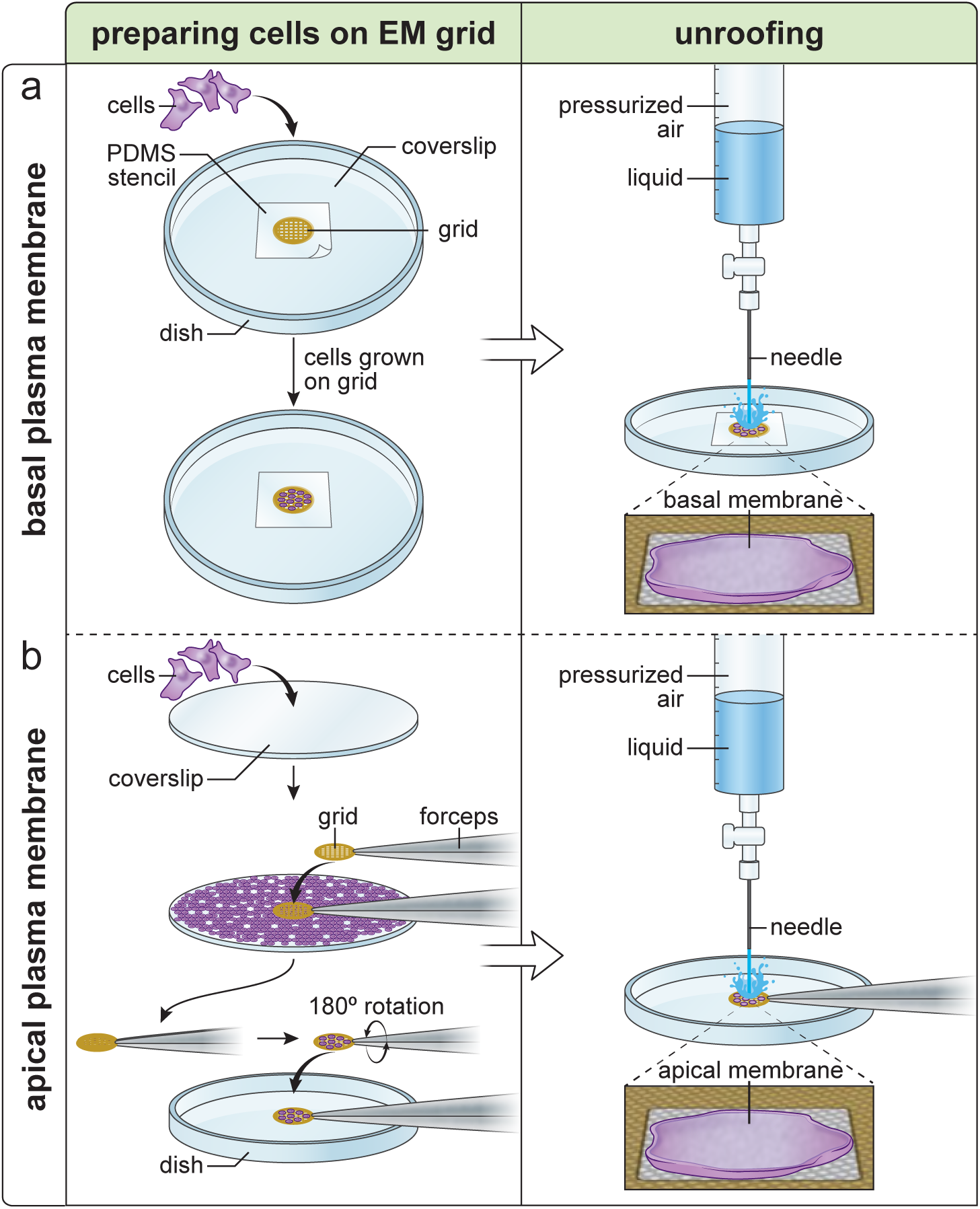
Generating isolated plasma membranes on EM grids. **a,** Diagrams showing the workflow for isolating basal plasma membranes. A plasma-cleaned grid is placed onto a coverslip and secured with a PDMS stencil. Cells are then seeded onto the grid and incubated overnight. To generate isolated basal plasma membranes, the grid is placed under a pressurized fluid-delivering device and the grid is sprayed with unroofing buffer to wash away the apical portions of cells with shearing force. **b,** Diagrams showing the workflow for isolating apical plasma membranes. Cells are first seeded onto a coverslip and incubated overnight. A plasma-cleaned, poly-L-lysine coated grid is then brought into contact with the coverslip to pick up cells. Pressurized fluid is then applied to the grid to wash away the basal portions of cells to generate isolated apical plasma membranes.

Apical plasma membrane preparations were performed by growing cells on a glass coverslip. To transfer the cells to an EM grid, the coverslip was rinsed in serum-free buffer and the carbon side of a poly-lysine-coated grid was lightly touched to the coverslip for 3-5 seconds and lifted (Fig. 1b). Cells transferred to the grid appeared largely intact as viewed by fluorescence, scanning electron microscopy, and FIB-milled cryoET (Fig. S2). The cells on the grid, now in an inverted orientation relative to their growth on the glass coverslips, were unroofed to generate isolated apical plasma membranes (Fig. 1b). Isolated basal and apical plasma membranes could then be used for blotting, vitrification, and imaging.

### Grid topography influences organelle distribution on membranes grown on grids

To characterize differences in the organization of cellular structures on grid-bound basal and apical HSC3 plasma membranes, we used platinum replica electron microscopy (PREM) (Fig. 2a-c). Apical membranes exhibited many filipodia. Both membrane types showed a range of actin filament organization, vesicles, caveolae, and clathrin-coated structures (Fig. S3). To quantify differences in the morphology of clathrin-coated structures within each preparation, we used semi-automated segmentation to identify three general classes: flat, dome, and spherical clathrin (Fig. 2d) [7]. The edge of the Quantifoil carbon film holes served as a reference to measure the distribution of clathrin-coated structures (Fig. 2e). Adherent basal plasma membranes exhibited more flat clathrin structures at or near the edge of the carbon film holes (Fig. 2f). Apical membranes exhibited no such preference (Fig. 2g). The addition of ultrathin carbon (2 nm) to the grids, which provides a carbon surface across the holes, lessened but did not eliminate this accumulation in basal membrane samples (Fig. S3). Overall, and as individual classes, the basal side contained more clathrin-coated structures (total projected area: 16.14 µm^2^ from 844 structures; 749.46 µm^2^ analyzed / # of structures per µm^2^: flat, 0.27 ± 0.19; dome, 0.35 ± 0.17; sphere, 0.47 ± 0.29) compared to the apical membrane (total projected area: 9.02 µm^2^ from 518 structures; 1036.44 µm^2^ analyzed / # of structures per µm^2^: flat, 0.15 ± 0.13; dome, 0.24 ± 0.15; sphere, 0.18 ± 0.13) (Fig. 2h).

**Figure 2.**
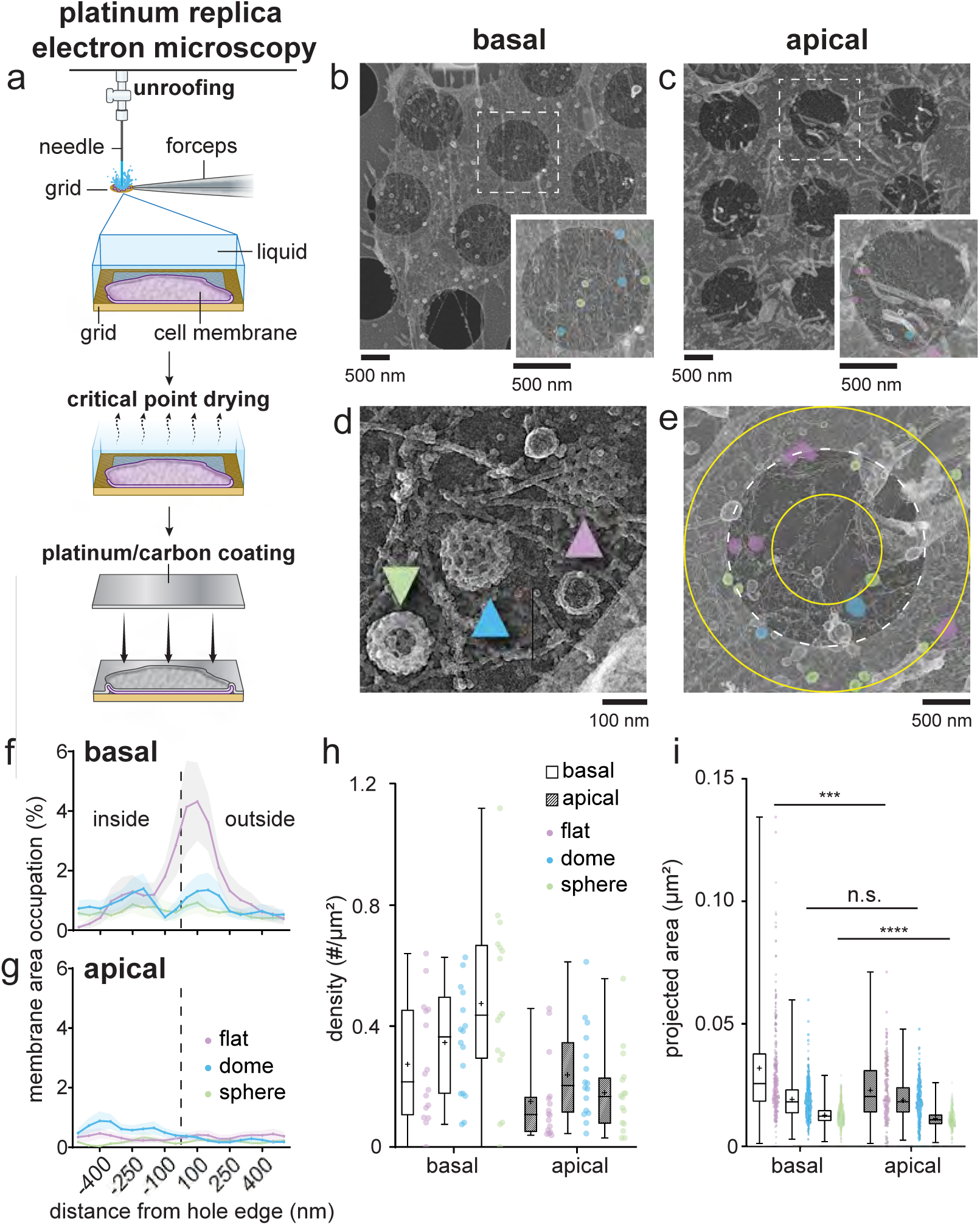
Evaluating isolated plasma membranes of HSC3 cells on EM grids with platinum replica electron microscopy. **a,** Cartoon showing the process from generating isolated plasma membranes on grids to producing platinum replicas of the isolated plasma membranes. **b,** An isolated HSC3 cell basal plasma membrane on an R2/1 Quantifoil grid. The inset shows an enlarged view of the white dash square area. Structural classes of clathrin are color-coded: Lilac = flat; processed cyan = dome; tea green = sphere. **c,** An isolated HSC3 cell apical plasma membrane on an R2/1 Quantifoil grid. The inset shows an enlarged view of the white dash square area. Color-coding is as in **b**. **d,** A close-up view showing the three classes of clathrin structures. The lilac-colored arrow points to a flat clathrin structure, with a visible clathrin lattice edge; the cyan-colored arrow points to a dome-shaped clathrin structure with an elevated lattice structure and a less well-defined lattice edge; the tea green-colored arrow points to a spherical clathrin structure with no visible clathrin lattice edge. **e,** The edge of the hole (white dash circle) is used as the reference point to evaluate the distribution of the different classes of clathrin-coated structures across the changing grid surface. Yellow circles inside and outside of the hole define the 500 nm range from the edge of the hole evaluated. The 1000 nm range (500 nm outside and 500 nm inside the hole edge) is divided into 10 bins of 50 nm (as plotted in **f,g**). Clathrin structures are color-coded as mentioned above. **f,g,** Comparison of the distribution of flat, dome, and sphere clathrin-coated structures with respect to the edge of the hole between basal **f** and apical **g** isolated plasma membranes. **h** and **i** are box and whisker plots, **h** comparing the density of different classes of clathrin structures between isolated basal and apical plasma membranes, and **i** of the projected area of individual clathrin structures grouped by structural class. On average, flat (***p = 0.0002) and sphere (****p < 0.0001) clathrin structures on basal membranes have a larger projected area when compared to their apical counterparts. For **h** and **i,** horizontal lines=median, plus signs=mean.

While the projected area of each class of clathrin structures varied (Fig. 2i), flat and sphere-shaped clathrin structures found in the basal membranes were larger (in µm^2^: flat, 0.032 ± 0.022; dome, 0.019 ± 0.008; sphere, 0.013 ± 0.003) than their apical counterparts (in µm^2^: flat, 0.023 ± 0.012; dome, 0.019 ± 0.008; sphere, 0.011 ± 0.004). Differences in macromolecular assemblies between the apical and basal membranes clearly demonstrate that when cells are grown on grids with complex topographies, the grid surface represents a unique structural challenge, and cellular objects may exhibit distinct distributions across the sample. Understanding these differences is key for future work on the structure and function of organelles on these heterogeneous substrates.

### Unroofing provides thin membrane samples suitable for cryoET

After unroofing, basal and apical membrane preparations of HSC3 cells were back-blotted, plunge-frozen, and observed with cryoET (Fig. 3a). With low or medium magnification imaging, membranes appeared light gray, only distinguishable by close inspection (Fig. 3b-c). Tomograms acquired from these membranes exhibited high contrast and were rich in membrane-bound organelles, actin, clathrin, intermediate filaments, and ribosomes (Fig. 3d-e). As sample thickness limits attainable resolution in cryoET [13, 14], the average thickness of each tomogram was measured. Basal tomogram thickness spanned from 78 to 196 nm and apical tomogram thickness spanned from 110 to 221 nm (Fig. 3f). The average thickness of isolated membrane tomograms used in this study was 163 ± 52 nm (mean ± stdv) (Fig. S4). Thus, unroofed samples have thicknesses rivaling the thinnest FIB-milled samples and provide high-contrast 3D views of a cellular environment.

**Figure 3.**
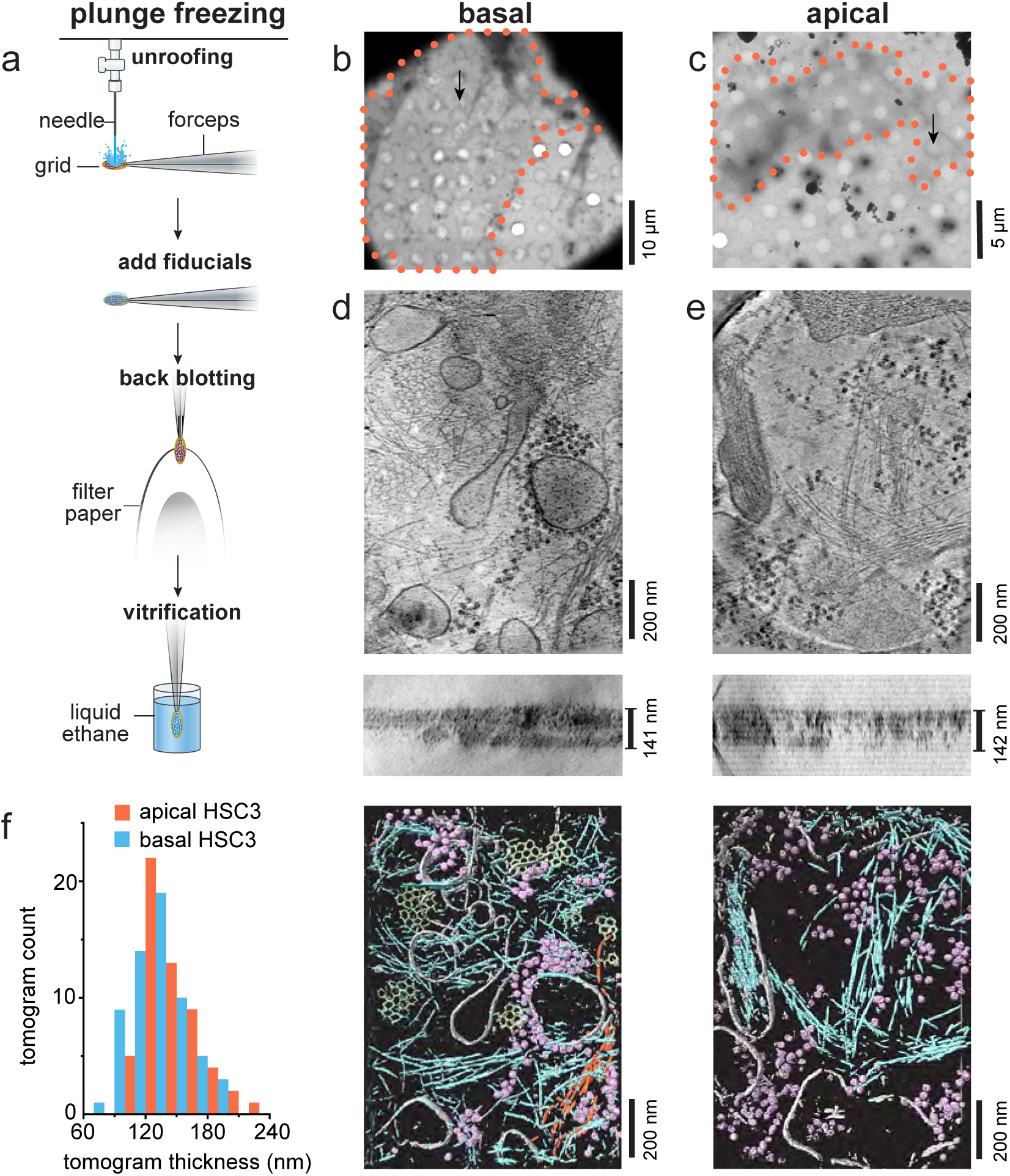
Unroofed cells provide 100-200 nm thick plasma membrane samples for cryoET. **a,** A diagram highlighting unroofing, addition of fiducial markers, back-blotting, and plunging in liquid ethane for vitrification. **b,** A 2250x magnification montage of a grid square containing an isolated HSC3 basal membrane (outlined with an orange dotted line). **c,** A 2250x magnification montage of a grid square containing an isolated apical HSC3 membrane (outlined with an orange dotted line). **d (top)** shows a minimum-intensity projection along the Z axis through 21 slices of a gauss-ian-smoothed bin8 tomogram acquired at the location of the arrow in **b**. **d (middle)** shows a minimum-intensity projection of 101 slices through the Y axis of the same tomogram with the measured thickness shown. **d (bottom)** shows a segmentation (mask guided isosurface) of the above tomogram: gray=membrane, purple=ribosomes, blue=actin, light green=clathrin, orange=intermediate filaments. **e,** Same as **d**, but for the apical membrane tomogram acquired at the position of the black arrow in **c**. **f,** Histogram of tomogram thicknesses of HSC3 basal and apical membranes imaged here. Napical=56, Nbasal=61, 2 grids represented for each. Scale bars are 10 μm, 5 μm for **b,c** respectively and 200 nm for **d,e.**

### High-resolution structural information is preserved in unroofed cells

Ribosomes were dense and recognizable in our tomograms. They were observed densely on or near internal membranes and dispersed at or above the plasma membrane. To assess the preservation of high-resolution structural information in vitrified unroofed samples, we performed subtomogram averaging (STA) on ribosomes in an apical preparation of HEK293 cells (Fig. 4a). 111 tomograms were used to refine 11,249 ribosomes to obtain a consensus structure of the full 80S ribosome at 7.5 Å nominal resolution (Fig. 4b-c, Fig. S5). Classification without alignment was performed and revealed two classes which resembled a pre-translocational rotated state and a non-rotated state (Fig. 4d, Fig. S5). The rotated state exhibited tRNA in the hybrid A/P, P/E sites (Fig. d, blue and pink respectively). The non-rotated state exhibited prominent density in the P state (Fig. d, orange). Separate focused classification at the peptide exit site revealed a coarse structure of a putative translocon and confirms the presence of plasma membrane proximal endoplasmic reticulum in many of our tomograms (Fig. 4e, Fig. S6). Together, these data show that particles found within unroofed membrane samples retain sub-nanometer structural information.

**Figure 4.**
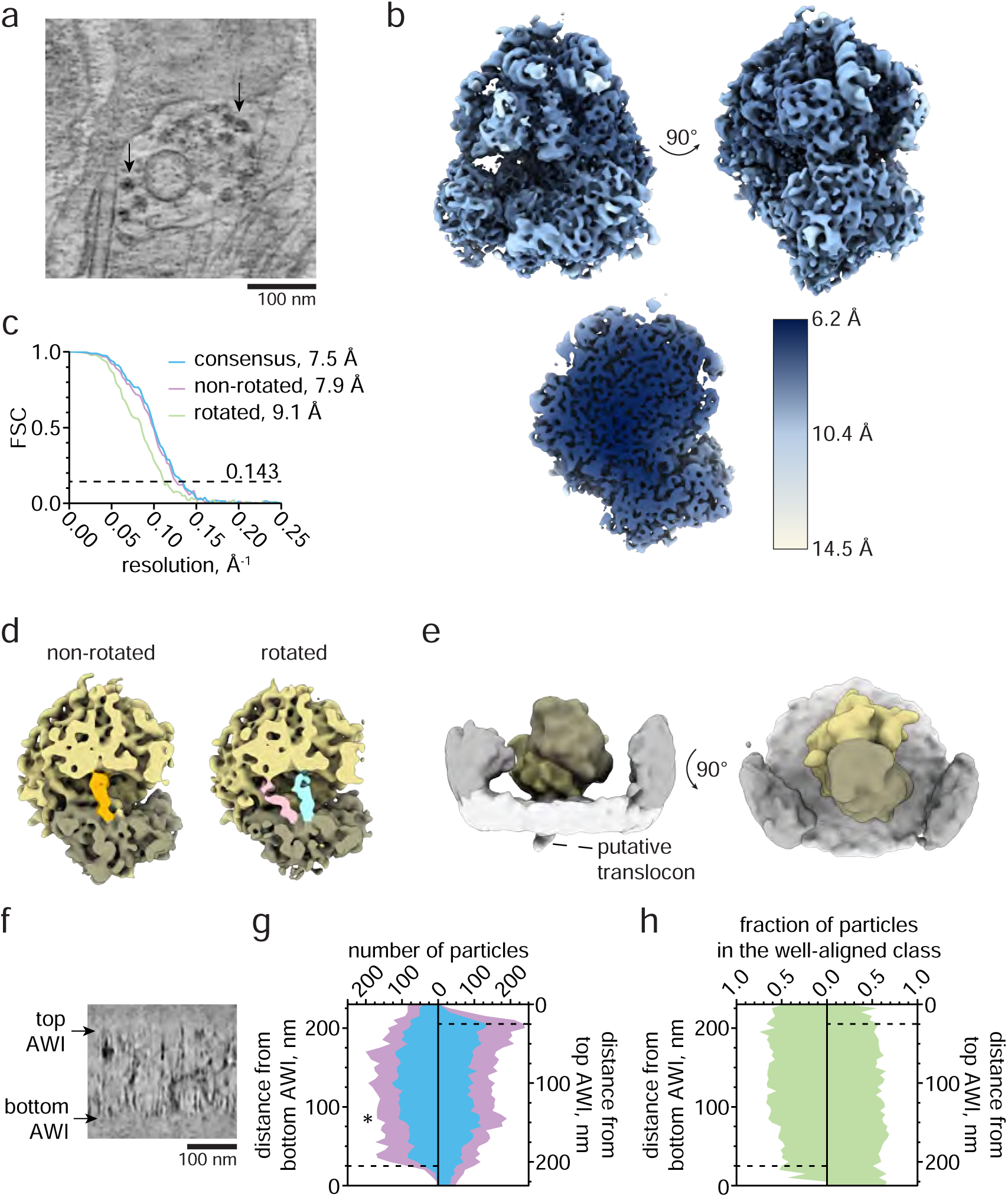
Subtomogram averaging and contextual analysis shows sub-nanometer detail is preserved in isolated plasma membranes. **a,** Projection of 21 z-slices from a tomogram of an isolated plasma membrane. 80S ribosomes (black arrows) are frequently found in unroofed HEK293 cells overexpressing dynamin-1(K44A). **b,** Rotated views (top) and a clipped view (bottom) of a consensus subtomogram average filtered according to the local resolution. **c,** Fourier shell correlation (FSC) profiles obtained from subtomogram averages. The nominal resolution is reported at FSC=0.143. **d,e,** Classification of the set of well-aligning particles obtained subtomogram averages of the 80S ribosome in non-rotated and rotated states. tRNA occupying the P, P/E, and A/P sites are indicated in orange, pink, and blue, respectively (**d**) and a subset of 446 membrane-bound ribosomes (two views, **e**). **f,** A view from a tomogram with the top and bottom air-water interfaces (AWIs) indicated (black arrows). **g,** Distances of putative ribosomes were measured from both AWIs. The number of particles falling within successive 5 nm bins from the bottom and top AWIs (left and right panels, respectively) is plotted for the set of particles obtained immediately following picking (cyan) and the set of well-aligning particles obtained from classification (purple). **h,** The fraction of particles found with particle picking that constitute the well-aligning class are plotted with respect to their distance to the bottom and top AWIs.

The air-water interface (AWI) is known to disrupt protein structures in *in vitro* cryo-samples [15]. Similarly, vitrified unroofed cells are bounded by two AWIs either proximal to the extracellular side of the plasma membrane (bottom) or distal to the intracellular side of the plasma membrane (top) (Fig. 4f). To examine how the AWI affected ribosome structure in unroofed samples, the AWI-particle distances were calculated from 111 tomograms (Fig. 4g). A peak in the particle distribution, indicating accumulation, occurred at 25-30 nm from the top AWI (Fig. 4g, asterisk). In contrast, the distribution measured from the bottom AWI gradually increased and plateaued at a distance of approximately 80 nm. This is likely due to the plasma membrane acting as a buffer between intracellular material and the AWI. The complete set of segmented particles (Fig. 4g, purple) was compared to the particles that grouped into well-aligned classes and therefore contributed to the final consensus map (Fig. 4g, cyan). The proportion of particles in the well-aligned class was used as an indicator of particle quality (Fig. 4h). On average, the percentage of particles selected for subsequent processing was mostly constant throughout the portion of the tomogram containing biological material. A rapid drop in the proportion of well-aligning ribosomes was observed ∼25 nm from both AWIs (dashed lines). This is consistent with the dimensions of ribosomes, which are ∼25 nm in size, being acutely damaged by AWI contact. We conclude that high resolution structural information for ribosomal complexes is retained in vitrified unroofed samples not in direct contact with the AWI.

### Correlative imaging facilitates tomogram acquisition of stalled endocytic events

To locate rare proteins or proteins of unknown structure, we implemented a CLEM protocol. The HEK293 cell line described above expresses, upon induction, a fluorescently labeled well-characterized dominant negative mutant of dynamin 1, Dyn1(K44A)-GFP, that blocks clathrin-mediated endocytosis [16–18]. Using this cell line, we tested our CLEM protocol to search for sites of arrested clathrin-mediated endocytosis. After apical unroofing, we added 500 nm red fluorescent poly-styrene beads which adhered to the grid, specifically in poly-L-lysine rich locations lacking plasma membrane. This image registration allowed us to precisely identify grid holes containing GFP fluorescence for tomogram acquisition (Fig. 5a-e). We acquired 198 tomograms on two grids and identified N=490 clathrin structures within the tomograms. We observed N=42 dynamin decorated tubules (9% of clathrin structures) where dynamin could be easily identified based on its characteristic polymer spiral (Fig. 5f) [16]. This is similar to the 10% previously reported in HeLa cells in resin sections [19]. Conversely, we commonly saw large arrested clathrin structures (N=219, 45% of clathrin structures) that resembled clathrin grape clusters (Fig. 5g) [20]. Though there was fluorescence at these sites, dynamin could not be identified in the tomograms, presumably because it was assembled in short spirals or other assemblies that were not readily distinguishable. The high density of arrested clathrin at these sites confirmed our ability to use CLEM to find sites where fluorescently labeled proteins were present. However, the inability to positively identify dynamin in each of these tomograms highlighted the need for a more precise EM-visible protein tag.

**Figure 5.**
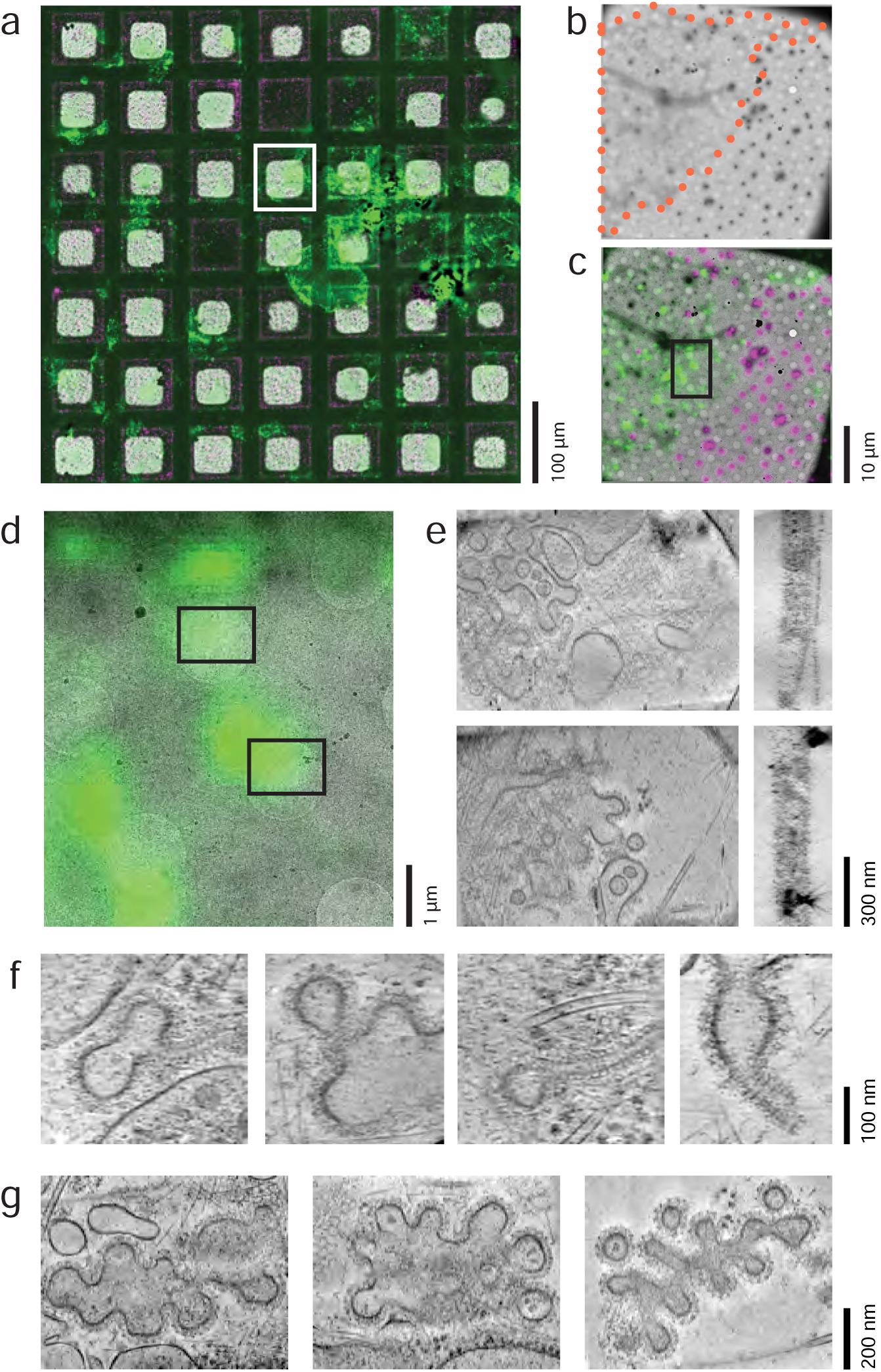
CLEM finds sites of arrested clathrin-mediated endocytosis (CME). **a,** Select portion of grid shown as a low magnification cryoEM image registered with cryo-fluorescence images of Dyn1(K44A)-GFP (green) and 500 nm fiducial markers (red). **b,** The grid square highlighted in white (**a**) is shown in a higher resolution map. Orange dots indicate the outline of the isolated plasma membrane. **c,** The fluorescence overlay is shown. **d,** The black box is shown enlarged. Black boxes indicate the location of tilt series acquisition for the tomograms shown in **e**. **e,** Examples of tomograms in XY (left) and XZ (right) **f,** Examples of arrested CME sites with Dynamin 1 (K44A) tubules. **g,** Examples of arrested CME sites with large clathrin-decorated clusters. All XY tomogram images are minimum intensity projections (mIPs) over 21 gauss-ian-smoothed XY slices while XZ images are mIPs of 101 XZ slices. Scale bars are **a,** 100 µm; **b,c,** 10 µm; **d,** 1 µm; **e,** 300 nm; **f,** 100 nm; **g,** 200 nm.

### Iron-free ferritin tagging identifies Hip1R in cryo-electron tomograms of unroofed cells

To identify specific proteins within a tomogram, we tested the ability of iron-free rapamycin-induced FerriTag [12] to label Hip1R, a 119 kDa, 50-60 nm long clathrin adaptor that makes a coiled-coil parallel homodimer linking the membrane to actin [21–25]. First, FerriTag recruitment to Hip1R was confirmed using fluorescence microscopy. Before rapamycin, HEK293 cells expressing Hip1R-GFP-FKBP and FerriTag exhibited diffuse red cytoplasmic fluorescence and green fluorescent puncta, typical of clathrin-associated proteins, in total internal reflection fluorescence microscopy (TIRF). With 200 nM rapamycin, the red fluorescence colocalizes with the green Hip1R puncta (Fig. 6a). This redistribution was rapid and visible within 30 seconds.

**Figure 6.**
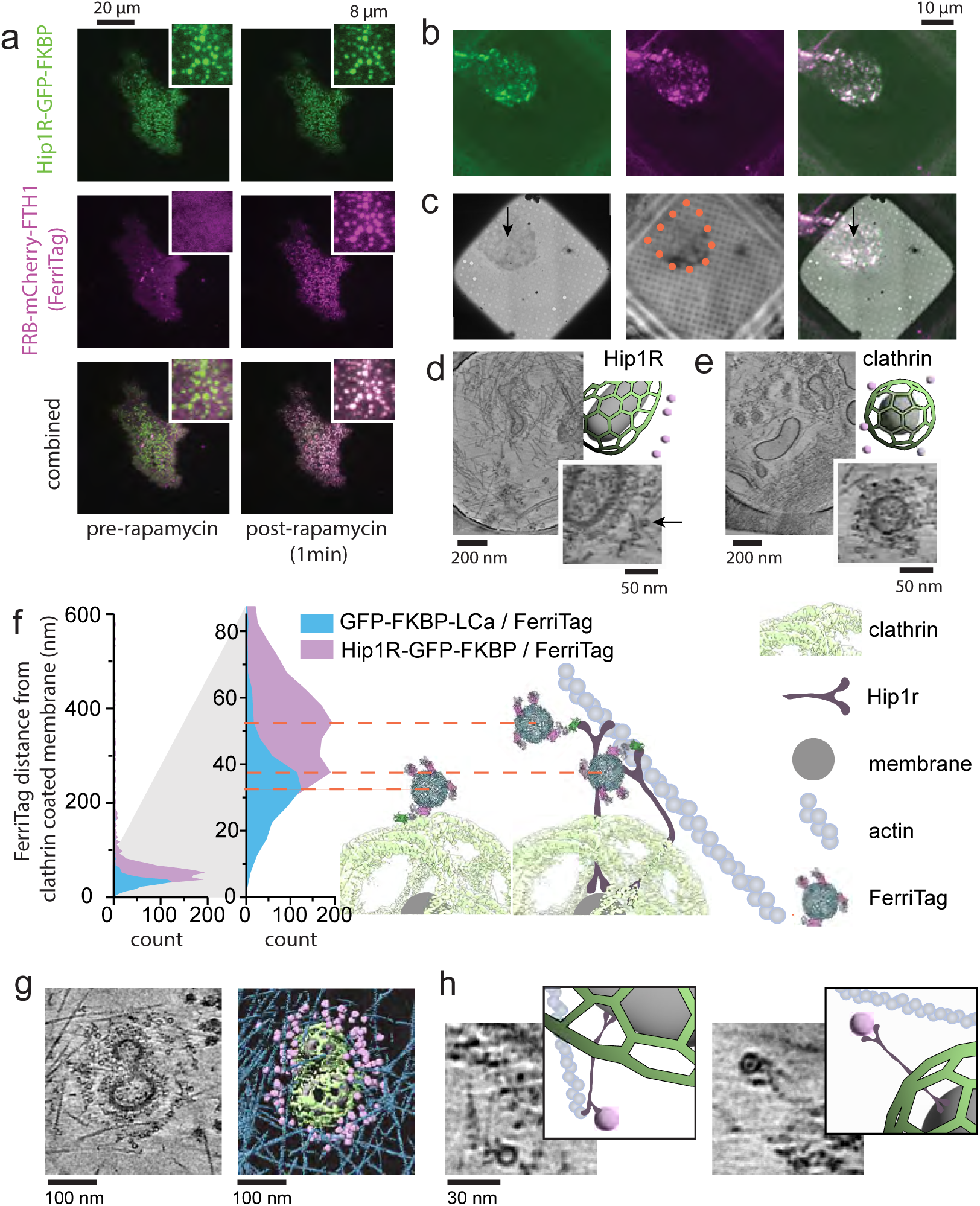
Iron-Free FerriTag is specific, efficient, and visible in cryoET. **a,** TIRF microscopy of a HEK293 cell expressing FerriTag (FRB-mCherry-FTH1, magenta, and FTL) and Hip1R-GFP-FKBP (green) shown before rapamycin addition (left) and 30-60 seconds after rapamycin addition (right). **b,** Cryo-fluorescence microscopy of HEK293 isolated plasma membrane on a grid expressing FerriTag (FRB-mCherry-FTH1, magenta, and FTL) and Hip1R-GFP-FKBP (green) with combined colors on the right. **c (left),** CryoEM montaged map of the same grid square. **c(middle),** Reflection image acquired on the cryo fluorescence microscope and used to register the images. **c(right),** Registered fluorescence and EM images combined. Orange dots = membrane outline, arrows = position of **d** acquisition. **d,** Tomogram of Hip1R/FerriTag labelling and an example of a prominent clathrin structure (right). Empty FerriTag structures are clearly visible as 12 nm circles (arrow). **e,** Selected tomogram from HEK293 cells with labeled FerriTag on GFP-FKBP-LCa shown and an example of a well-defined clathrin structure (right). For **d** and **e**, a cartoon at the top right depicts the location of the membrane (gray), clathrin (green), and FerriTag (purple). **f,** A histogram of the FerriTag distance from clathrin coated membrane (shown at 0-600 nm and 0-80 nm) for clathrin light chain (GFP-FKBP-LCa) and Hip1R (Hip1R-GFP-FKBP). Molecular models of Ferritin (PDB-1fha), mCherry (PDB-2h5q), FRB/FKBP (PDB-3fap), and GFP (PDB-5wwk) and EM density of the clathrin cage (emdb-21608) at scale with the zoomed-in histogram in combination with cartoons of Hip1R, actin, and membrane are shown as putative models explaining the data to the left. The dashed orange line guides the eye from the data peaks to the center of the FerriTag model. **g,** An example of FerriTag Hip1R labeling and surrounding a clathrin structure. Segmentation of membrane (gray), clathrin (green), FerriTag (purple) and actin (blue) are shown to the right. **h,** Close-up examples showing Hip1R density adjacent to FerriTag. Cartoons of membrane (light gray), clathrin (green), Hip1R (dark gray), FerriTag (purple), and actin (blue) are shown to the right to aid image interpretation. Tomogram images in d,e are minimum intensity projections (mIPs) of 21 Gaussian-smoothed XY slices. Tomogram images in **g,h** are mIPs of 10 Gaussian-smoothed slices. Scale bars are **a,** 20 μm, 8 μm inset full-width; **b,c,** 10 μm; **d,e,** 200 nm, 50 nm inset; **g,** 100 nm; **h,** 30 nm.

Next, we used the Hip1R-FerriTag system to examine empty ferritin cages in cryoET. After 2-minute incubation in 200 nM rapamycin, we unroofed cells and observed the apical membranes in cryo-fluorescence and cryoET. At sites with GFP and mCherry fluorescence, tomograms exhibited distinct 12-nm hollow spheres surrounding clathrin lattices (Fig. 6b-d). These spheres were prominently visible and could be automatically identified using a trained convolutional neural network (EMAN2). Distributions of Hip1R/FerriTag (as in Fig. 6d) were compared to that of FerriTag coupled to the N-terminus of clathrin light chain A (GFP-FKBP-LCa; as in Fig. 6e), a subunit of the heterohexamer triskelia that sits at the membrane-distal portion of the ∼25 nm thick clathrin membrane coat. For both Hip1R and clathrin, the data exhibit a large peak within 100 nm of the clathrin-coated membrane and a very low background (clathrin, grid #7-8, 91 tomograms, 81% within 100 nm, N_<100_=633, N_tot_=784; Hip1R, grid #9, 68 tomograms, 81% within 100 nm, N_<100_=1638, N_tot_=2057)(Fig. 6f). Of the data within 100 nm, FerriTag is found 35 ± 15 nm away from the membrane for GFP-FKBP-LCa and is 50 ± 17 nm away for Hip1R. The Hip1R peak plateaus between 38 and 52 nm, consistent with Hip1R radially projecting from the clathrin mesh, or tilted at an angle (Fig. 6f, model). Segmentation and enlarged tomogram slices show that several Hip1R can bind to a single actin fiber and are directly observed in extended and angled states (Fig. 6g-h). Together, these data demonstrate that iron-free ferritin is an efficient, expressible, chemically-induced EM and light visible probe for cryoET in unroofed samples.

## Discussion

We present a correlative cryoET pipeline for imaging proteins at the plasma membrane. An air-pressure-driven syringe unroofing technique provides a quick and simple method for preparing isolated grid-bound apical or basal membranes from mammalian cells. With PREM, we show that growing cells on a Quantifoil grid alters the organization of the basal membrane cortex, specifically quantified for clathrin. The thickness and content of the material in vitrified unroofed samples were found suitable for obtaining sub-nm protein structure with STA. Correlative cryo-fluorescence and iron-free FerriTag complement cell unroofing to identify membrane-associated proteins of interest. These methods facilitate visual proteomics of the eukaryotic cell plasma membrane.

Our PREM analysis of grid-bound membranes was performed on a single human cell line (HSC3). It is expected that contents between the apical and basal membranes will differ in a cell-type dependent manner. However, evidence supports the idea that flat clathrin will likely concentrate at the edge of carbon holes in basal membranes (Fig. 2f) in most adherent cell types. Specifically, clathrin has been reported to concentrate over sites of high substrate curvature in MDA-MB-231 breast cancer cells [26]. Our observation of flat clathrin lattices at edges is consistent with the longer average clathrin-mediated endocytosis (CME) lifetimes observed at high substrate curvature [26]. Flat clathrin lattices in HSC3 cells, rat myotubes, and many other cell lines have been directly linked to substrate adhesion with integrins [27–29]. Similarly, adhesive clathrin structures aggregate at high-curvature collagen-pinching membrane ridges during cell migration [30]. Thus, flat clathrin structures, observed in many cell types [7], are expected to exhibit adherent patches at the edge of carbon holes.

The AWI is well-known to cause protein adsorption, preferred orientation, and damage in single particle cryoEM [31]. Here, unroofing exposes cellular material to an AWI. Notably, membrane-bound protein complexes are blocked from the bottom AWI by the membrane and most are kept from the top AWI due to their physical association with the plasma membrane. Ribosome positions in our data confirm that proteins can accumulate as the top AWI shrinks the buffer volume during blotting (Fig. 4g). Protein complexes in contact with the AWI likely experience damage (Fig. 4h). The AWI damage seems to be more acute than radiation damage that occurs during FIB-milling [32]. While the STA performed here on ribosomes achieves similar resolution to other FIB-milled ribosomal RELION 4.0-based STA [33], higher resolution has been obtained with FIB-milled samples with other STA software [34–36]. However, tomograms of unroofed cells exhibit particular advantages for STA efforts. The decreased cytosolic crowding makes protein shape and position distinct and, coupled with the lack of FIB radiation damage, enhances potential for high-resolution STA.

To identify specific proteins in the tomograms, we used an empty FerriTag to label both Hip1R and clathrin light chain. Clathrin light chain, in particular, is an integral component of a dynamic and interlaced protein cage where bulky tags are potentially problematic. Still, the brief chemically-induced tagging required for FerriTag facilitates close labelling while allowing for an intact (Fig. 6e) and functioning lattice [12]. We have previously visualized the distribution of Hip1R around clathrin structures in HeLa cells using correlative super-resolution localization fluorescence microscopy and PREM [37]. With this lower-resolution CLEM technique, the averaged fluorescence signal was consistent with Hip1R being distributed throughout the clathrin lattice with the N-terminus splayed radially outward in curved structures. The cryoET presented here is consistent with previous results but at much higher resolution. When Hip1R/FerriTag was previously visualized in resin sections [12], the average distance of FerriTag from the coated membrane was 30 nm, 20 nm smaller than what was measured here. It is possible that 70 nm sections result in a 2D projection of Hip1R that is angled toward or away from the plane of reference, making the distances appear shorter. A recent cryoET study of CME actin forces, showed putative Hip1R densities in the intact human skin melanoma cell line, SK-MEL-2 [38]. These ∼50 nm-long densities contain two arms at either end, cross the clathrin lattice, and are consistent with what we observed here. FerriTag helps us to confidently identify these long, thin and relatively dim densities as Hip1R, showcasing the utility of the techniques presented.

The potential of cryoET and cryo-fluorescence is expanding, and its impact on cell biology and medicine will depend on wider access. Greater accessibility requires more rapid and available sample preparation techniques. Here, we show that the preparation of vitrified unroofed cells is a powerful addition to the fast-evolving cryoET toolbox. These methods enable imaging of the many important plasma membrane proteins involved in the function of cells and diseases.

## Methods

### Cell lines

The human tongue squamous cell carcinoma cell line, HSC3, used here is genome edited to express epidermal growth factor receptor (EGFR)-GFP (at 40-50% of EGFR) and was a kind gift from Dr. Alexander Sorkin [39]. It was maintained in DMEM (Thermo-Fisher 11995065) supplemented with 10% FBS, with or without 50 mg/mL streptomycin – 50 U/mL penicillin (Thermo-Fisher 15070063). HEK293 WT cells were purchased from ATCC (CRL-3216) and maintained in Eagle’s minimum essential medium (EMEM, Fisher Scientific 502382632) supplemented with 10% fetal bovine serum (FBS). For FerriTag experiments, HEK293 cells were transfected with *FTL* (Ferritin light chain, Addgene plasmid # 100750), *FRB-mCherry-FTH1* (Ferritin Heavy Chain, Addgene Plasmid #100749), and one of either *HIP1R-GFP-FKBP* (Addgene Plasmid #100752) or *GFP-FKBP-Lca* (Addgene Plasmid #59353) [12] using the Lonza Nucleofector and kit V (Lonza #VCA-1003) immediately prior to coverslip seeding. 200 nM rapamycin was added to the cells for the indicated times to induce FKBP/FRB heterodimerization. The inducible HEK293 Dynamin1(K44A)-GFP cell line was generated as follows. The cDNA coding for human Dynamin1-K44A-GFP (Addgene #34681) was inserted into *pcDNA5-FRT/TO* to generate *pcDNA/TO/Dyn1-K44A-GFP* using the In-Fusion cloning kit (Takara), and the following primers: ATGGTGAGCAAGGGCGAGG, ACGCTAGAGTCCGGAGGC, TCCGGACTCTAGCGTCCTGCCATGGGCAACCGC, GCCCTTGCTCACCATGGTGGCGACCGGTGGATCC. Proper insertion was confirmed by full sequencing (Plasmidsaurus). To generate inducible cells expressing Dynamin1(K44A)-GFP, parental Flp-In T-Rex HEK293 cells (Thermo Fisher, R78007) were co-transfected with a 1:5 mass ratio of *pcDNA/TO/Dyn1-K44A-GFP* and the Flp recombinase expression plasmid, *pOG44* (Thermo Fisher, V600520) using Lipofectamine 3000 (Invitrogen, L3000-015) following manufacturer’s instructions. 24 h after transfection, cells were subjected to selection with 5 μg/mL blasticidin and 100 μg/mL hygromycin B while grown in DMEM supplemented with 10% tetracycline-free FBS until the formation of cells grown in isolated colonies [40]. Single colonies were isolated using cloning glass cylinders (Sigma-Aldrich, C1059) and the cells were further expanded. We confirmed the presence of Dynamin1-K44A-GFP after inducing the transgene expression with 1 μg/mL doxycycline hyclate for 18 h. We observed the presence of diffraction-limited GFP puncta across the plasma membrane of 100% of the cells using TIRF, as shown before [41]. After selection, Dyn1-K44A-GFP Trex HEK293 cells were maintained in EMEM with 10% tetracycline-free FBS and 5 μg/mL blasticidin. For experiments, induction was performed with 100 ng/mL doxycycline hyclate overnight. All cells were maintained in sterile conditions at 37°C in 5% CO_2_. They were confirmed mycoplasma free and ATCC-authenticated.

### Growing HSC3 cells on EM grids

EM grids (R2/1 Au 300 for PREM with no carbon; Au 300 R2/2, 2 nm carbon for cryoET and PREM with carbon) were first plasma cleaned for 30 seconds using a PELCO easiGlow (Ted Pella Inc.) and then secured to glass coverslips (1 grid/coverslip) using PDMS stencils (Nanoscale Labs 8052901) with the carbon-coated side facing up and placed into a 35 mm dish. EM grids were placed into the cell culture hood for subsequent steps starting with 20 minutes of UV light sterilization. Next, grids were coated with fibronectin (diluted with autoclaved H_2_O to reach a final concentration of 125 ng/µL, Sigma-Aldrich, F1141) for 20 minutes. Coated grids were washed with sterile H_2_O before cell seeding with an average of 15k cells (contained in a 20 µL droplet) per EM grid. Grids were transferred to an incubator at 37 °C with 5% CO_2_ for 1 hour to allow the cells to settle. After 1 hour, 4 mL growth media was added to each dish and the dishes were returned to the incubator for overnight incubation.

### Growing cells on coverslips

HSC3 cells were seeded onto 25 mm diameter rat tail collagen I-coated coverslips (Neuvitro Corporation, GG-25-1.5-collagen) with an average of 400k cells per coverslip. Cells were then incubated overnight at 37 °C with 5% CO_2_. HEK293 cells were seeded onto Fibronectin-coated coverslips (NeuVitro Corporation, GG-25-1.5-Fibronectin) and grown overnight before use.

### HSC3 serum-starving

HSC3 cells used for PREM and basal cryoET were serum-starved prior to unroofing. After overnight incubation, cells were washed with PBS at 37 °C and then synchronized by incubating the cells in starvation buffer (DMEM supplemented with 1% v/v Glutamax and 10 mM HEPES) at 37 °C for 1 hour followed by incubation in starvation buffer supplemented with 0.1% w/v BSA at 4 °C for 40 minutes.

### Cell unroofing

For generating isolated basal plasma membranes, an EM grid with cells was first washed in PBS at room temperature and then placed under an air pressure-driven fluid delivery device (Fig. S1) (ALA Scientific Instruments Inc.). The unroofing buffer (2% paraformaldehyde in stabilization buffer [30 mM HEPES, 70 mM KCl, 5 mM MgCl_2_, 3 mM EGTA, pH 7.4]) is applied to the EM grid at between 0.7 and 0.8 bar above atmospheric pressure for 1-2 seconds. Next, the EM grid is either washed with stabilization buffer and placed into a Leica Microsystems plunge freezer for vitrification using liquid ethane at -180 °C or placed into 2% PFA for subsequent processing for generating platinum replica.

To prepare for isolating apical plasma membranes, EM grids (Quantifoil R2/1 Au 300, Q3100AR1 for PREM, per Table 1 for cryoET) were plasma cleaned for 30 seconds and coated with 0.01% poly-L-lysine (Sigma-Aldrich, P4832) for 20-60 minutes. Cells grown on coverslips were washed with stabilization buffer and placed in stabilization buffer during the transfer of cells from a coverslip to an EM grid. Coated EM grids were washed with H_2_O or stabilization buffer and then brought into contact with cells on a coverslip for 3-5 seconds to pick up cells off the surface. The cells on the EM grid were then unroofed and processed as described above.

**Table 1.**
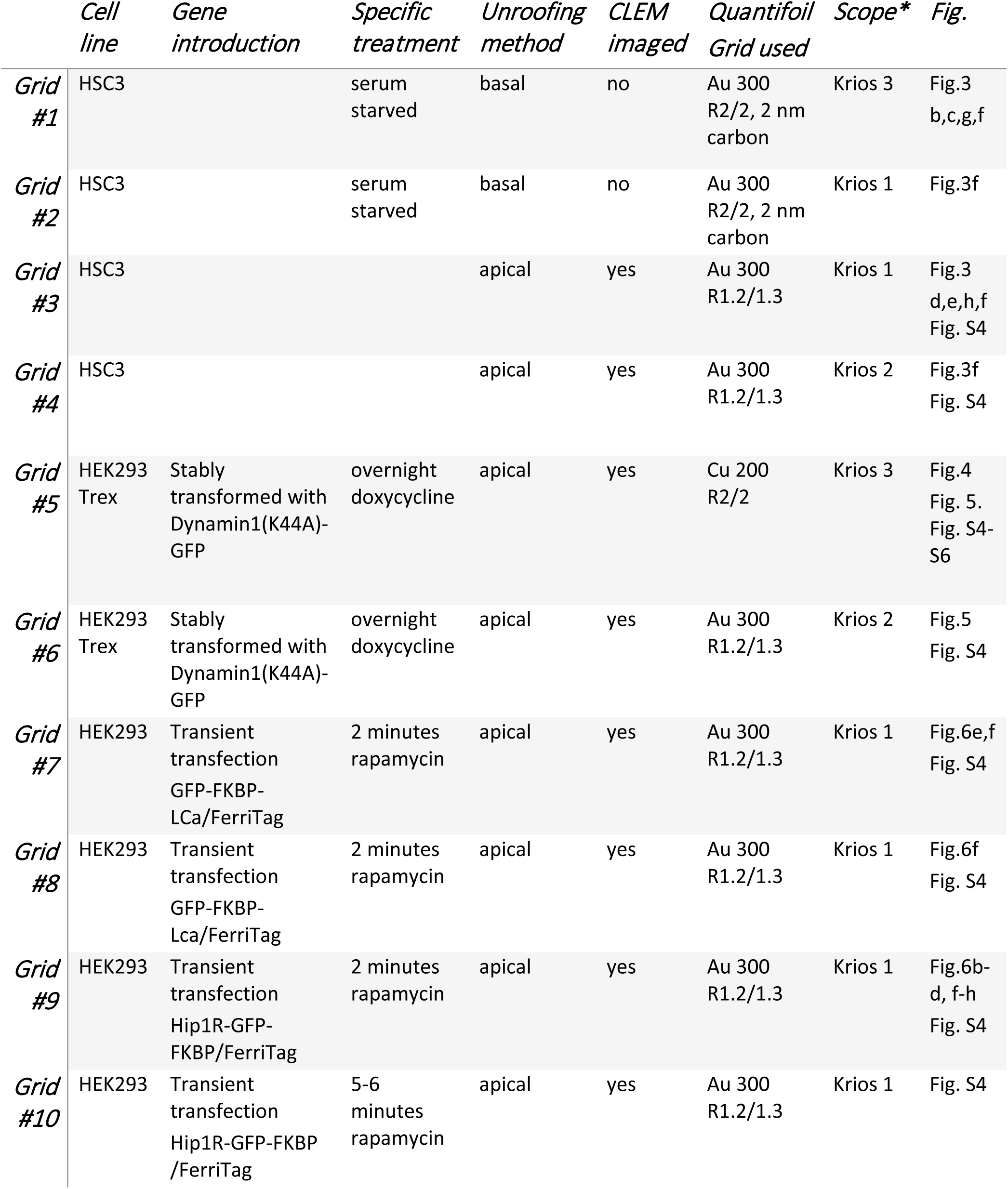
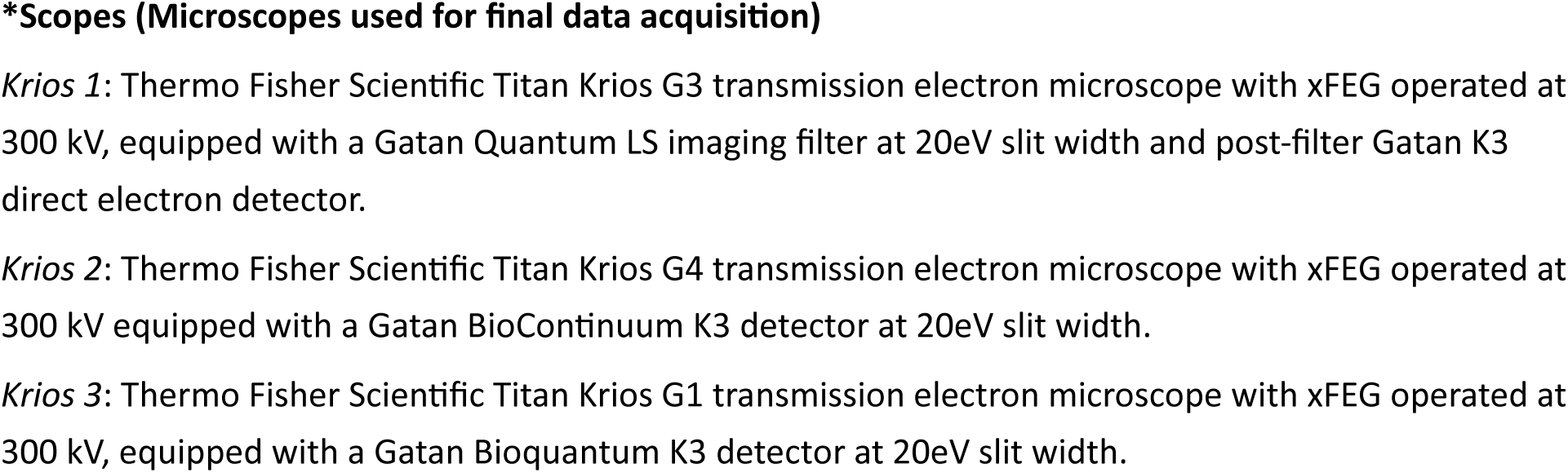
CryoET Sample summary.

### Vitrification

EM grids were washed in stabilization buffer and lightly blotted from the side, prior to adding 4 µL of fiducial solution (10 nm BSA gold tracer, Electron Microscopy Sciences, 25486; 1:1 with stabilization buffer for grids). The fiducial solution also included 500 nm carboxylate modified red fluospheres (Invitrogen F8812) for grids #5 (50 µg/mL), grids #3,4,6 (25 µg/mL). Plunge freezing was performed on a Leica GP plunge freezer. The chamber was conditioned to 25 °C at 80% humidity for grids #1-2. The chamber was conditioned to 15 °C and 95% humidity for all other grids. Grids were blotted from behind for 3 seconds and then plunged into liquid ethane at -180 °C. Vitrified grids were transferred to liquid nitrogen for storage.

### Platinum replica electron microscopy

The generation of platinum replica on an EM grid follows the steps as previously described [27] with slight modifications. Unroofed cells on EM grids were placed into 2% paraformaldehyde for 20 minutes and then transferred into 2% glutaraldehyde at 4 °C overnight. Following overnight fixation, EM grids were incubated in 0.1% tannic acid (in H_2_O) for 20 minutes at 4 °C, washed with 4 °C ddH_2_O, and then incubated in 0.1% uranyl acetate (in H_2_O) for 20 minutes at 4 °C. Grids were washed extensively with ddH_2_O post uranyl acetate and underwent dehydration stepwise in ethanol (15%, 30%, 50%, 70%, 80%, 90%, 100% x3) at room temperature using 4 minutes per step. EM grids were then critical point dried using an Autosamdri-815 (Tousimis). Grids were then transferred to a Leica EM ACE 900 for coating with a platinum thickness target of 3 nm (17°, 40 rpm, 110W) and a backing carbon thickness target of 2.5 nm (90°, 40 rpm, 150 W). Coated EM grids were imaged directly (biological material still present under metal coat) using a 120 kV FEI Tecnai T12 equipped with a Rio 9 CMOS camera (Gatan). Montages were collected at 5.87 Å/pixel resolution using SerialEM [42]. Montage blending was done using the IMOD software suite [43]. We evaluated plasma membrane-grid surface interaction by segmenting and quantifying different classes of clathrin structures. Segmentation of three classes of clathrin structures— flat (no visible curvature), domed (curved lattice with a visible edge), and sphere (curved beyond a hemisphere and no visible lattice edge)—was carried out using the deep learning model Mask R-CNN [44, 45]. Model training was done on 113 manually generated segmentation masks [7]. Output segmentation masks were manually reviewed, using custom built widgets for the image analysis platform Napari [46]. Masks of the whole plasma membrane area segmented, holes in the Quantifoil, and each class of clathrin structure were combined in FIJI [47]. The composite masks were then processed with MATLAB to analyze the density, projected area, and spatial organization of clathrin structures in relation to the edge of each hole in the Quantifoil carbon film for each class of clathrin structures. Projected areas were collected by counting pixels per each mask. The densities were calculated by counting the number of each type of clathrin structure mask in a given area analyzed. To evaluate the spatial organization of clathrin structures with respect to the edge of a Quantifoil hole, we first assign a centroid to the shape of each clathrin structure mask. Concentric rings, each 50 nm thick, were used to analyze a range from 500 nm inside and outside of the hole edge (20 rings; 10 inside, 10 outside). Membrane area occupation was defined as the percentage of pixels in the 50 nm-thick ring that were occupied by the segmented clathrin mask from that particular class. Density was defined as the number of structures per area analyzed. Projected area was the area of each clathrin structure in the segmented masks. Results from MATLAB were output to an Excel file for subsequent data organization. Statistical comparisons were performed using the Mann-Whitney test (one-sided). A *p* value of <0.05 was considered statistically significant. Data visualization and statistical tests were done using Prism (GraphPad). For basal membranes on Quantifoil, analysis was performed on N=16 membranes, 203 flat, 237 domed, 404 spherical clathrin structures from 2 grids. For apical membranes, analysis was performed on N=16 membranes, 139 flat, 206 domed, 173 spherical clathrin structures from 1 grid. For basal membranes on Quantifoil with 2 nm carbon, analysis was performed on N=10 membranes, 226 flat, 184 domed, 273 spherical structures from 1 grid.

### Cryo-fluorescence microscopy

Cryo-fluorescence imaging was performed on a CryoCLEM Thunder Imager (Leica Microsystems) where indicated in Table 1 (Grids 3-10). Full grids were imaged with a GFP filter cube (Exc:470/40,dichroic 495, Em:525/50 Leica #11504164), a Y3 filter cube (Exc:545/25,dichroic 565, Em:605/70 Leica #11504169), and a reflector cube (BF-LP425, Leica #11505287). Final images used were single-slice extended focus projections of 35 µm thick (1 µm increments) montaged stacks (130 nm/px). Registration of the fluorescent data onto the corresponding EM atlas allowed affine transformation and was accomplished within MATLAB to target areas of interest for tomogram acquisition.

### Cryo-electron microscopy

Vitrified unroofed samples were screened by acquiring low magnification grid atlases on a 200kV Thermo Fisher Glacios. Electron tomography was carried out on one of three 300kV Thermo Fisher Titan Krios as indicated in Table 1. Low magnification grid maps were acquired at 135x magnification. After coarse fluorescence image registration, grid square maps were acquired at 2250x magnification. Fluorescence image registration was finely adjusted to new grid square maps to identify tomogram positions of interest. Tilt series were collected dose-symmetrically at 42000x magnification with a grouping of 3 using SerialEM [42] at a resolution of 1.08 Å/pixel (K3 at super-resolution mode) and 4-5 subframes per tilt from -50° to +50° at 2° increment and an evenly distributed dose targeted at 120 electrons/Å^2^.

### Tomogram reconstruction

Tomograms were reconstructed for general purpose visual analysis separately than they were reconstructed for STA. For general purpose, subframes were aligned at bin 8 with IMOD [48] “alignframes”. For STA, subframes were aligned at bin 1. Tomogram reconstruction was performed using IMOD batch processing “batchruntomo”. For general purpose, the batchruntomo directives included automatic X-ray removal, no additional binning (maintaining the bin 8 from the frame alignment), used 10 fiducials for global (no local) alignment, fiducial erasing with noise using 12 px extra diameter, and a back-projection tomogram reconstruction with a SIRT-like filter equivalent to 6 iterations. The batchruntomo directives included automatic X-ray removal, a target of 100 fiducials for global (no local) alignment, binning by 10 after stack alignment (resulting in bin10 final tomograms, 10.8 Å/px), fiducial erasing with noise using 4 px extra diameter and three expanded circle iterations, and a back-projection tomogram reconstruction with a SIRT-like filter equivalent to 10 iterations. The CTF was estimated per-tilt in IMOD, then processed with IsoNet [49] including CTF deconvolution, denoising, and modeling the missing wedge to improve the performance of automated particle picking in EMAN2 [50, 51].

### Sample thickness evaluation

Using the general purpose tomogram reconstructions at bin8, each tomogram was sampled at nine equally spaced XY positions throughout the tomogram. At each location, a region of 200x200 voxels in XY were cropped out, Gaussian-smoothed, and observed with a minimum intensity projection along the Y-axis in the XZ plane. Using this view, the thickness was the manually determined distance between the two AWIs. For each tomogram, the average of these nine values was reported as the tomogram thickness.

### FerriTag analysis

FerriTag was automatically segmented using a trained convolutional neural network in EMAN2 [51]. The output was thresholded at a 1.4 confidence level to create a 3D mask. A watershed function was used to separate nearby tags. Regions with fewer than 100 voxels were removed from the mask. The centroid of each region was used as the FerriTag position. Tags that were within 25 nm of the AWIs and tags within 75 nm of the tomogram XY edge were not used for analysis. The AWI was determined with manual segmentation and interpolation. The membrane under clathrin coats were manually segmented in IMOD and converted to 3D masks using the imodmop function. The nearest distance between each FerriTag position and the resulting clathrin-coated membrane masks was determined using the bwdist function in Matlab.

### CryoET segmentation

For Figs. 3g-h and 6g, actin, ribosomes, membrane, intermediate filaments, and FerriTag were all initially segmented with EMAN2 to create a 3D mask. These masks were manually corrected as needed to remove errant identifications. For clathrin, manual segmentation and masking was performed (IMOD, ImageJ). The masks were used to isolate signal within the tomogram for isosurface rendering. The final displayed segmentations are mask-guided isosurface renderings. Membrain [52] was used to generate the membrane segmentation in Fig. S6b.

### Subtomogram averaging and classification

Particle positions were imported into Dynamo [53] and subtomograms were extracted at a binning factor of 10 to generate an initial reference map. Pseudo-subtomograms were extracted and refined at a binning factor of 8 (8.66 Å/px) followed by two rounds of classification with a soft spherical 320 Å diameter mask where junk particles were removed. The remaining 43,996 particles were visualized using the ArtiaX [54] plugin for ChimeraX [55] in each tomogram. A population of ribosomes were located at the underside of the carbon film, possibly due to positively charged poly-lysine-coated carbon film attracting negatively-charged ribosomes from the cytoplasm during unroofing. These film-adhered ribosomes were discarded from analysis. Refinement in RELION [56] was performed on the remaining 23,619 particles at a binning factor of 4 (4.33 Å/px) using a soft mask around the entire complex. Further, refinement was performed at binning 2 (2.165 Å/px) with a mask focusing on the LSU, followed by classification without alignment (K=10). One class, containing 11,288 particles, was chosen, based on the rlnAccuracyRotations, rlnAccuracyTranslationsAngst, and rlnEstimatedResolution parameters. Structures of non-rotated and rotated ribosomes were obtained by classification (K=2) without alignment of the consensus particle set using a mask focused on the SSU. In a separate classification (K=12), a single class of membrane-bound ribosomes (446 particles) was obtained from the consensus particle set without alignment using a mask focusing on the peptide exit tunnel and the surrounding region (Fig. S5). Three refinement cycles of CTF and frame alignment parameters with refinement of the consensus particle set were performed. Final maps were obtained after unbinned refinement. Maps were locally filtered and sharpened in RELION 4.0.

### Ribosome distribution analysis

Tomograms processed with IsoNet, at 10.825 Å/px, were examined in 3dmod and points were manually defined to model the top and bottom AWIs in tomograms of unroofed cells. The top and bottom interfaces were determined with model points placed based on visual inspection of the tomograms, fiducial positions, and ice. A custom Python script was used to interpolate the picked model points for each interface and calculate distances from each particle to the interpolated surfaces.

### Live TIRF microscopy

HEK293 cells transfected with *FTL*, *FRB-mCherry-FTH1*, and *HIP1R-GFP-FKBP* were grown on fibronectin-coated coverslips and imaged with total internal reflection fluorescence (TIRF) microscopy using a Nikon Eclipse TI inverted fluorescence microscope with a 100× apoTIRF 1.49 NA objective and 488-nm, and 561-nm excitation lasers. Each color was imaged with 100 ms exposure every 30 seconds. The sample was kept in growth media at 37 °C and 5% CO_2_ during imaging. 200 nM rapamycin was added between frames of the movie.

## Data and code availability

Binned tomograms are being made available on the Chan Zuckerberg Initiative CryoET Data Portal. Ribosome structures are being made available as EMD-44921, EMD-44909, and EMD-44922. Code used for subtomogram averaging is available at https://github.com/dmichalak/sta-pipeline. Code for image correlation is available at https://github.com/KASochacki/clemposo.

All other materials or code is available upon request.

## Acknowledgements

We thank the NIH Intramural CryoEM Consortium (NICE), and the NIH Multi-Institute Cryo-Electron Microscopy Facility (MICEF) for use of equipment, data acquisition support, and data management support; specifically, Rick Huang of NICE, and Huaibin Wang, Bertram Canagarajah, and Ulrich Baxa of MICEF. We thank Alexander Sorkin (University of Pittsburgh, USA) for the generous gift of the HSC3/EGFR-GFP cell line; Jiamin Liu of the NIH Advanced Imaging and Microscopy Resource (AIM) for help training mask R-CNN for platinum replica segmentation; NIH HPC Biowulf cluster for computational resources; Naoko Mizuno for discussions during project initiation; Ethan Tyler of NIH Medical Arts for creating Fig. 1. This work was funded by Chan Zuckerberg Initiative, Visual Proteomics Imaging program. JWT is supported by the Intramural Research Program (IRP) of NHLBI, NIH. JEH is supported by the IRP of NIDDK, NIH.

## Author Contributions

WWS, DJM, KAS functioned as the core team to acquire, discuss, analyze data, and write manuscript. KAS was team leader. WWS performed/analyzed PREM and cryoET grids 1-2. DJM performed STA on ribosomes. KAS performed/analyzed cryoET grids 3-10. PK segmented tomograms. MAA and M-PS performed/supported plasmid/cell line preparations. AMA provided python script support for PREM segmentation. KAS, JWT, JEH guided experiments with regular feedback. WWS, DJM, KAS, JEH, and JWT edited manuscript.

