## SupplementaryFigsS1-S6 for "Cryo-electron tomography pipeline for plasma membranes"

### Figure S1.

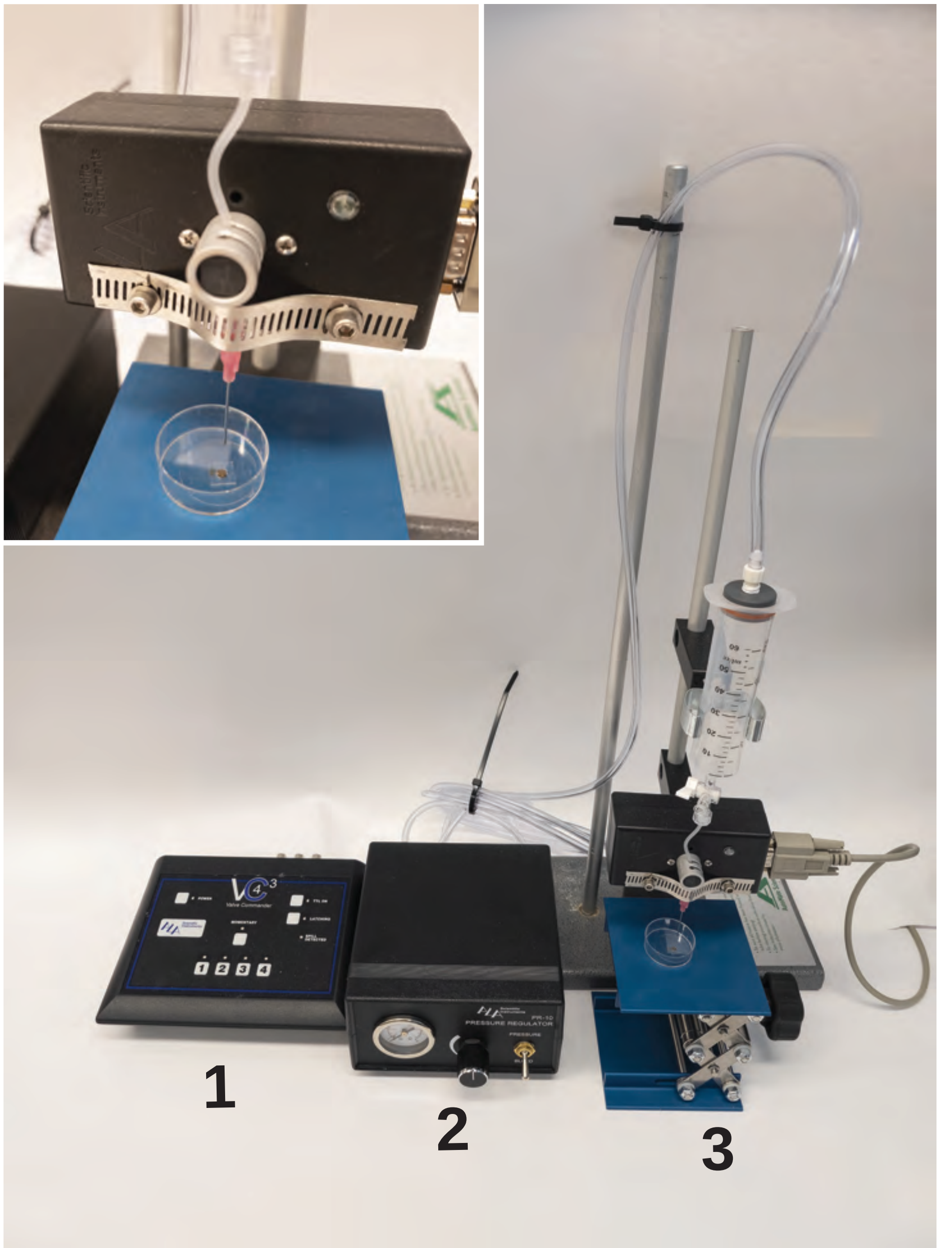


**Figure S1. Cell unroofing setup.** The unroofing system consists of valve control (1), pressure control (2), and a syringe (3) that applies a pressurized stream onto a cell-containing EM grid. The inset shows a close-up view of the EM grid-stencil-coverslip setup.

### Figure S2.

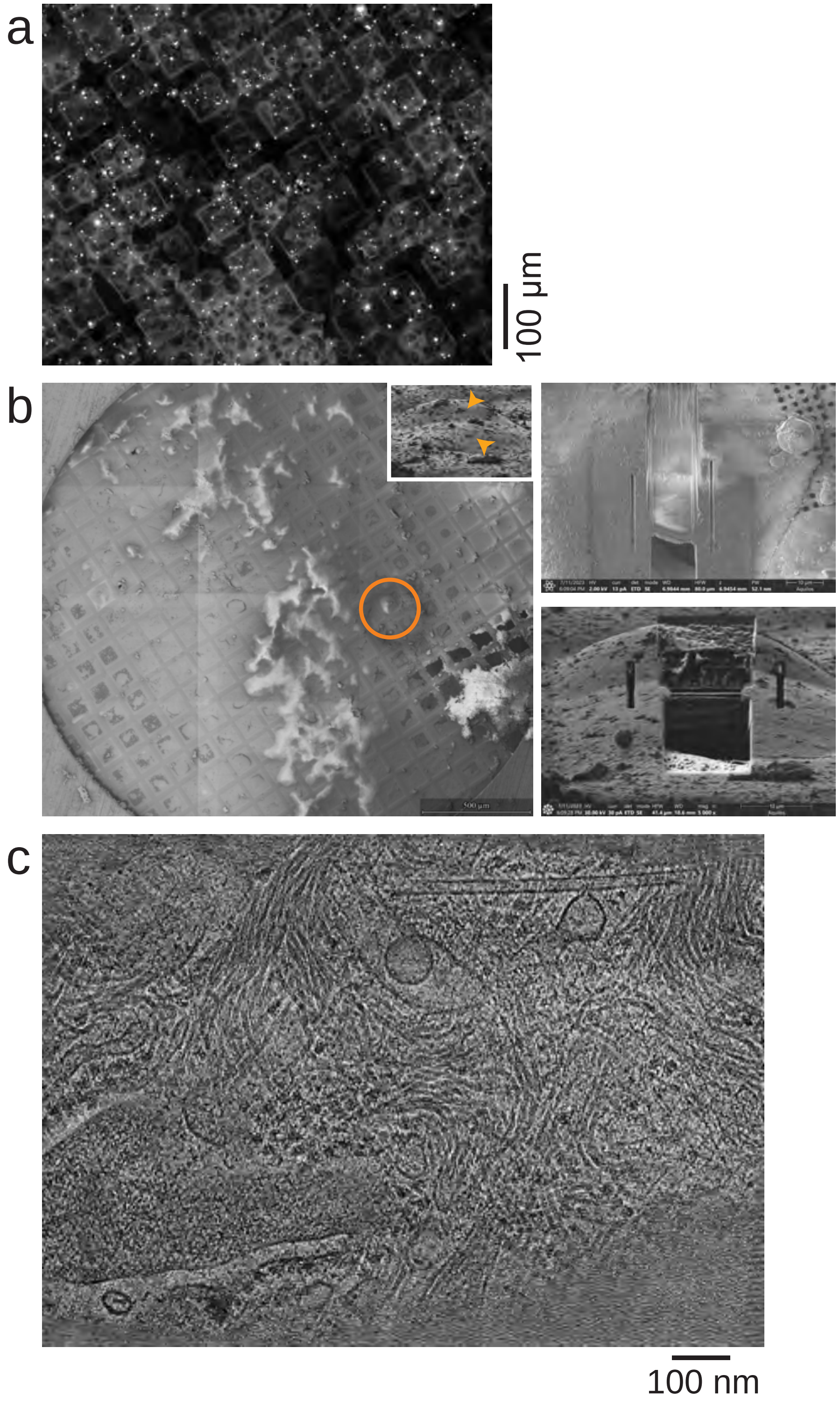


**Figure S2. Evaluating cells on EM grids and FIB milling of vitrified HSC3 cells. (a)** A fluorescent image from Leica CLEM showing intact HEK293 cells transfected with Dynamin-K44A-GFP on a poly-L-lysine coated EM grid. **(b)** A series of images showing the FIB-milling workflow. HSC3 cells were picked up by a poly-L-lysine coated EM grid and plunge frozen. Left panel, backscattered electron view and ion view (inset) showing the overview of a vitrified EM grid with picked-up HSC3 cells and a close-up view (orange circle) of two cells in a grid hole (orange arrows). Right panels show the backscattered electron (right-top) and ion (right-bottom) view of a finely milled lamella. **(c)** A tomographic projection from a tomogram collected from a lamella. A variety of features are preserved in an HSC3 cell including microtubules, ribosomes, intermediate filaments, and membrane organelles.

#### Figure S3.

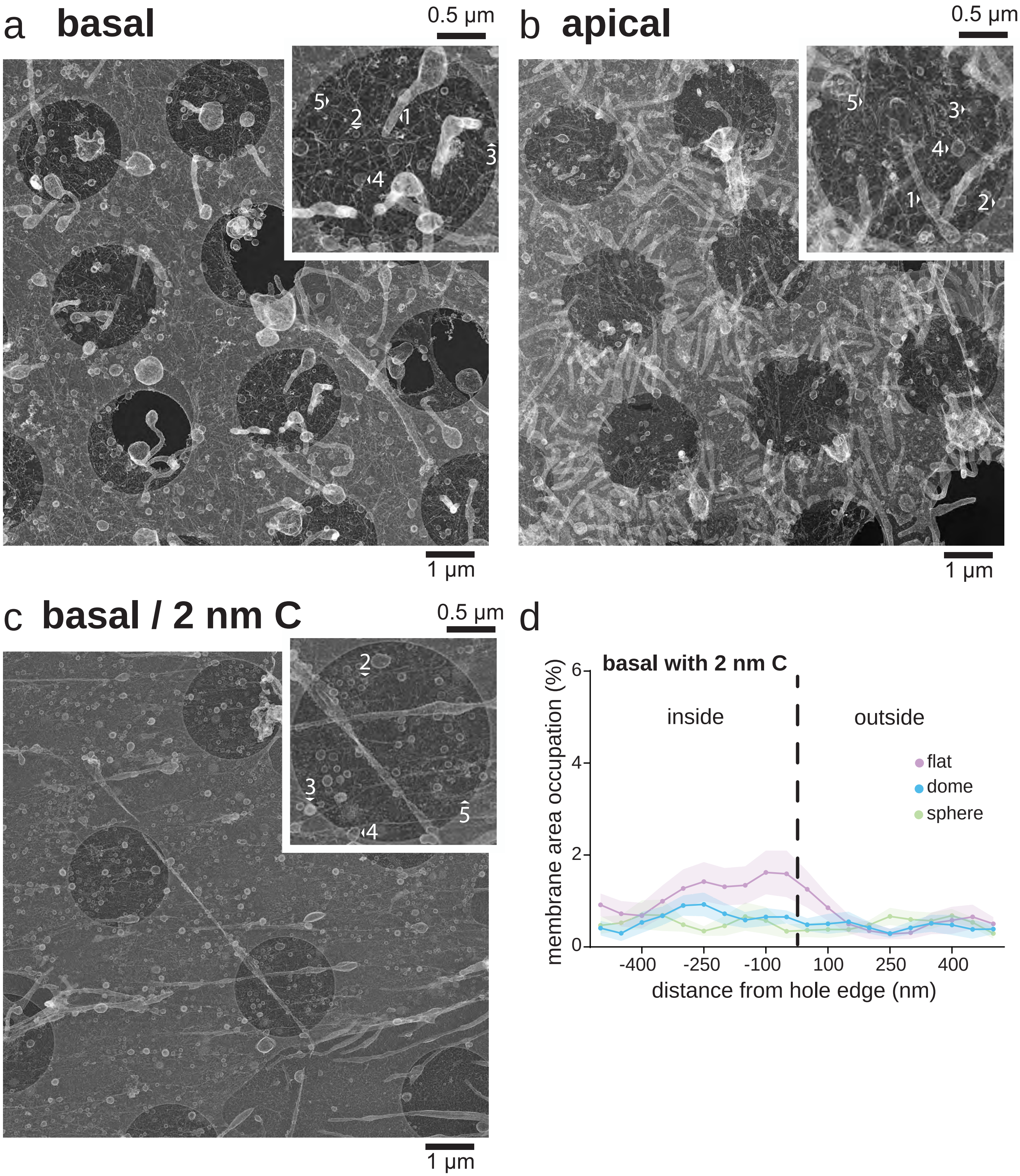


**Figure S3. Platinum replica electron microscopy of different HSC3 cell plasma membrane isolation preparations. (a)** An HSC3 basal plasma membrane on a Quantifoil R2/1 Au grid with enlarged crop out. 1=filipodia, 2=caveola, 3=clathrin, 4=vesicle, 5=actin. **(b)** An HSC3 apical plasma membrane on a Quantifoil R2/1 Au grid with enlarged crop out. **(c)** An HSC3 basal plasma membrane on a Quantifoil R2/2 +2 nm carbon Au grid with enlarged crop out. **(d)** The distribution of flat, dome, and sphere clathrin-coated structures for isolated basal plasma membranes from HSC3 cells grown on Quantifoil R2/2 +2 nm carbon Au grids. All images are tiled and stitched.

### Figure S4.

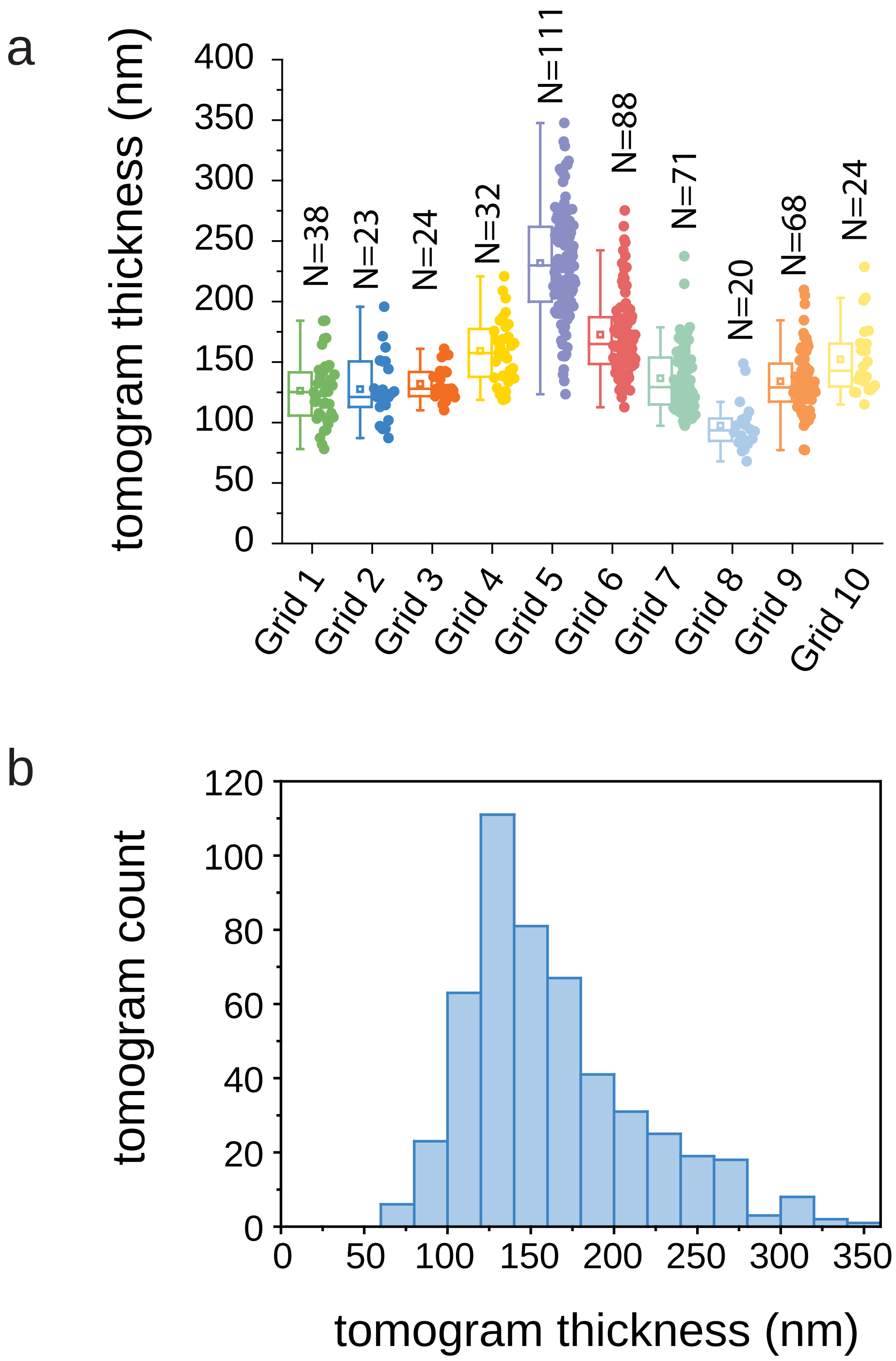


**Figure S4. Tomogram thickness. a,** The thickness of each cryo-tomogram used in this study grouped by grid. Grid numbers correlate with the numbers listed in Table 1. Boxes indicate interquartile range with median (line), mean (square), and outlier range (whisker, coeff. 1.5). **b,** The same data as in a but combined into a single histogram.

### Figure S5.

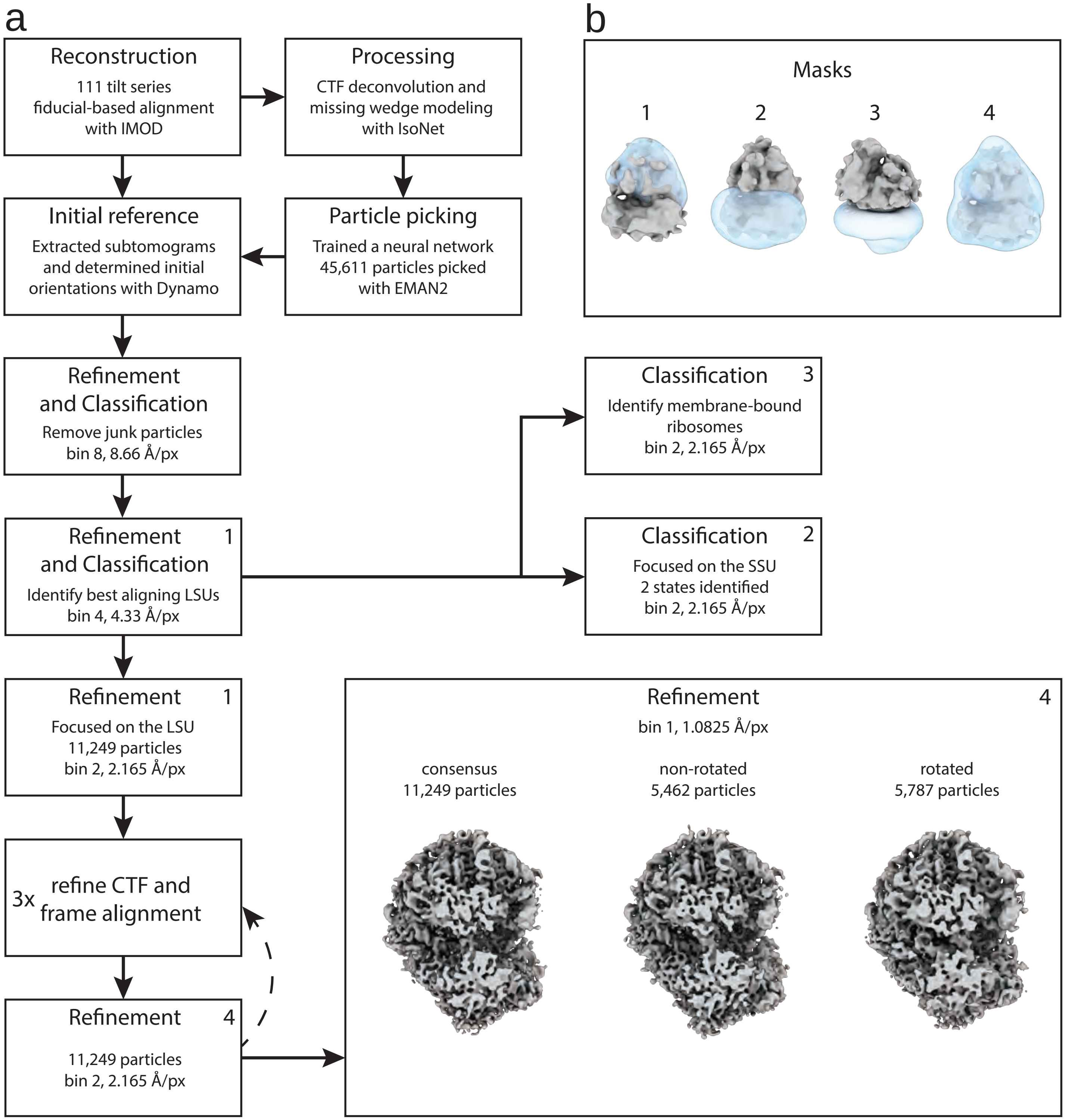


**Figure S5. Visual representation of the subtomogram averaging workflow. a,** Particles were subject to multiple rounds of refinement and classification to remove junk particles and poorly aligning ribosomes. The rotated and non-rotated particle populations defined by the SSU-focused classification were used in subsequent refinements. **b,** Several masks were used in different processing steps focused on the ribosomal LSU (1), SSU (2), peptide exit tunnel and membrane region (3), and a full mask (4). A number in the upper right corner for a processing step indicates the mask applied.

### Figure S6.

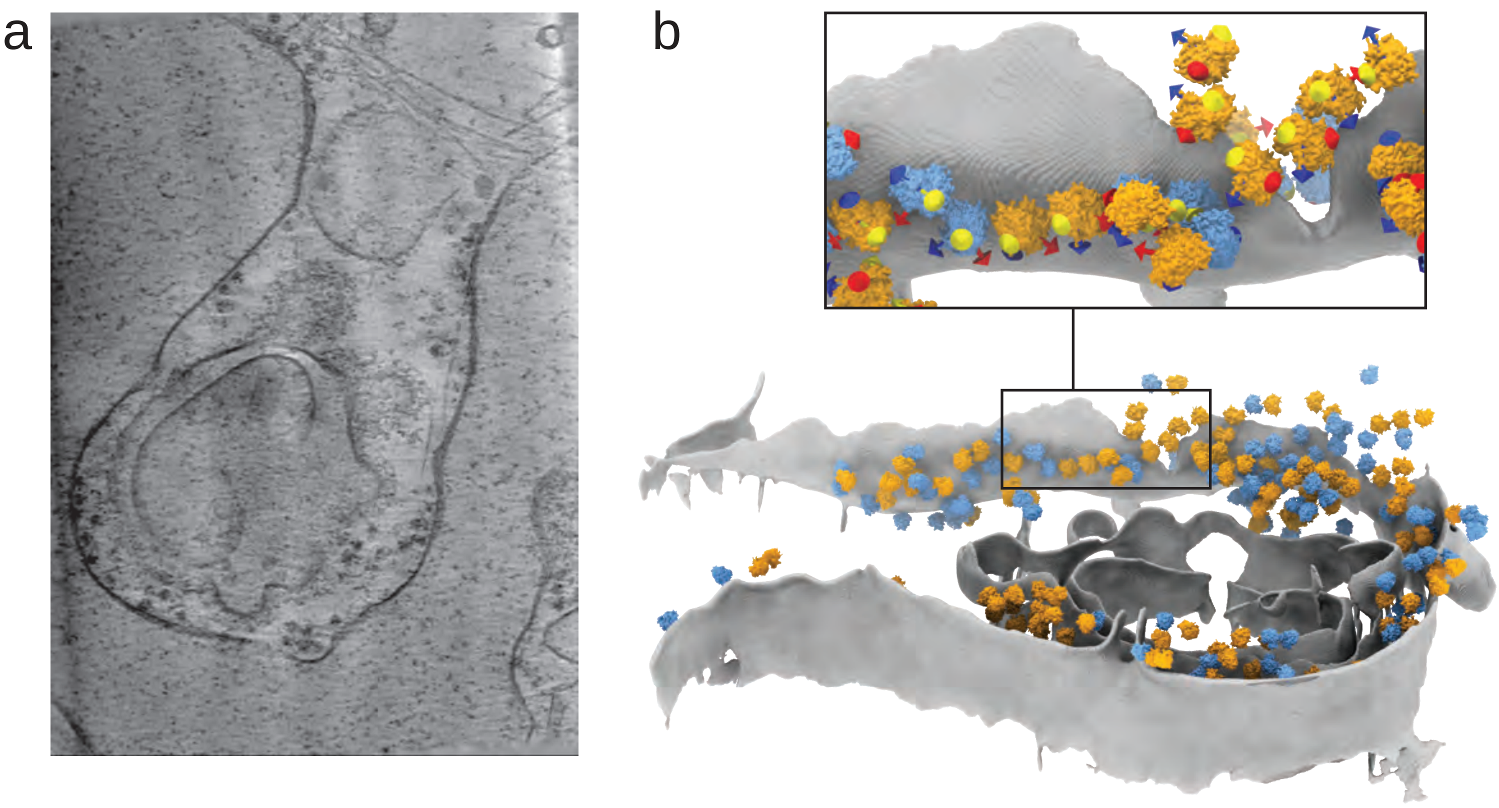


**Figure S6. ER-bound polysome in unroofed membrane tomogram**. **a**, Z-axis projection of 21 slices from a tomogram containing ER-bound ribosomes. **b**, Segmented ER membrane (gray) is shown with bound ribosomes. Averages from rotated (blue) and non-rotated (orange) classes are superimposed on particle positions from the membrane-bound particle set identified from classification. Similarly aligned orientation axes of neighboring ribosomes indicate a polysome (inset).
